# Generative modelling of the thalamo-cortical circuit mechanisms underlying the neurophysiological effects of ketamine

**DOI:** 10.1101/688044

**Authors:** Alexander D Shaw, Suresh D Muthukumaraswamy, Neeraj Saxena, Rachael L Sumner, Natalie Adams, Rosalyn J Moran, Krish D Singh

## Abstract

Cortical recordings of task-induced oscillations following subanaesthetic ketamine administration demonstrate alterations in amplitude, including increases at high-frequencies (gamma) and reductions at low frequencies (theta, alpha). To investigate the population-level interactions underlying these changes, we implemented a thalamo-cortical model (TCM) capable of recapitulating broadband spectral responses. Compared with an existing cortex-only 4-population model, Bayesian Model Selection preferred the TCM. The model was able to accurately and significantly recapitulate ketamine-induced reductions in alpha amplitude and increases in gamma amplitude. Parameter analysis revealed no change in receptor time-constants but significant increases in select synaptic connectivity with ketamine. Significantly increased connections included both AMPA and NMDA mediated connections from layer 2/3 superficial pyramidal cells to inhibitory interneurons and both GABA_A_ and NMDA mediated within-population gain control of layer 5 pyramidal cells. These results support the use of extended generative models for explaining oscillatory data and provide *in silico* support for ketamine’s ability to alter local coupling mediated by NMDA, AMPA and GABA-A.

## Introduction

The spectral composition of magnetoencephalographic (MEG) signals derived from visual cortex reflect a complex system of neuronal interactions, revealed as frequency-specific oscillations (Brunel and Wang, 2003; Buzsáki and Wang, 2012) and power law-like phenomena. Individual differences in common metrics, such as the peak frequency and amplitude within specific frequency windows, can predict behaviours (Ward, 2003), disease states (Hansenne, 2006; Herrmann and Demiralp, 2005; Nishida et al., 2011; Özerdem et al., 2008; Perry et al., 2014; Shaw et al., 2019; Tan et al., 2013), and are sensitive to experimental manipulations, including pharmacological manipulations (Campbell et al., 2014; Magazzini et al., 2016; Muthukumaraswamy, 2014; Muthukumaraswamy et al., 2016; Saxena et al., 2013; Shaw et al., 2015).

The frequency content of visual cortex typically contains prominent oscillations in theta / alpha (4 – 7, 8 – 13 Hz), through beta (13 - 30 Hz) up into the gamma (30+ Hz) range (Bastos et al., 2014; Clayton et al., 2017; Gray and Singer, 1989; Hall et al., 2005; Perry et al., 2013; Schmiedt et al., 2014; Xing et al., 2012). Subanaesthetic doses of ketamine, an uncompetitive NMDA receptor antagonist, have been shown to modulate both ends of this spectrum during task, with decreases in the amplitude of lower frequencies (Hong et al., 2010; Lazarewicz et al., 2010) and increases in the amplitude of higher frequencies (Hong et al., 2010; Shaw et al., 2015).

While the laminar and network generators of frequency-specific oscillations remain equivocal, prevailing theories propose distinct, dominant laminar generators for oscillations of different frequencies. One particular proposal is that high frequencies tend to be formed by current densities in supragranular layers, while lower frequencies appear to predominate in deeper regions of cortex, around layer 5/6 (Clayton et al., 2017; Maier et al., 2010) or in thalamo-cortical loops (Halgren et al., 2017; Lorincz et al., 2009). Under this model, the noted effects of ketamine on both low and high frequency bands suggest altered neuronal connectivity across much of the cortical laminae, and possibly in thalamo-cortical connectivity.

Computational modelling is an approach based on the use of basic mathematical descriptions of neuronal units, such as neural mass models, to explore neurophysiological phenomena. Methods such as Dynamic Causal Modelling (DCM) (Friston et al., 2003; Kiebel et al., 2008; Moran et al., 2007, 2011b) provide frameworks for generating typical and atypical empirical features in electrophysiological time series (e.g. (Adams et al., 2020; Cooray et al., 2015; Gilbert et al., 2016; Shaw et al., 2019)). In DCM, the subsequent fitting (inverting) of parameterised neuronal models to empirical data features, acquired under different pharmacological or task states, permits an *in-silico* assay of usually unobservable neuronal states such as synaptic connectivity between cell populations, or the decay times of specific receptor types. This approach has received construct and face validations (Moran et al., 2011a; Phillips et al., 2016).

We implemented a conductance-based thalamo-cortical circuit, with realistic intrinsic dynamics, to assay changes in both cortical and thalamo-cortical population connectivity and receptor dynamics underlying observed spectral changes induced by a subanaesthetic dose of ketamine, as recorded with magnetoencephalography (MEG). Our motivation to derive a neurophysiologically-informed, extended thalamo-cortical model was driven by observations that more reduced models lack the dimensionality to recapitulate broadband (i.e. theta through to gamma) spectra without help from a shaped noise model (Shaw et al., 2017). Furthermore, several theories propose a role for thalamo-cortical communication in the generation and pace-making of lower frequency oscillations (Halgren et al., 2017; Lumer, 1997). Hence, we built a model which contains the necessary cytoarchitectural apparatus to generate a range of oscillations (i.e. superficial, granular, deep and thalamic components). We expect that this model will provide a superior fit as it is able to better recapitulate the data. This is demonstrated when we compared our model to the default DCM NMDA cortical model (cmm_NMDA.m) using Bayesian Model Selection.

We hypothesised that our novel model would outperform the cortical model and that alterations in NMDA and possibly AMPA receptor mediated activity would be identified under ketamine, in line with previous modelling reports (Gilbert et al., 2018; Muthukumaraswamy et al., 2015). However, the direction of effects of NMDA and AMPA mediated connectivity in networked cortical models is equivocal, with both increases (Gilbert et al., 2018) and decreases (Muthukumaraswamy et al., 2015) reported. We also sought to explore whether ketamine altered local connectivity measures mediated by other receptors, given the complex binding profile of ketamine (Zorumski et al., 2016).

## Materials and Methods

### Sample characteristics

Details of the sample and procedures of this study have been reported previously, see Shaw 2 (Shaw et al., 2015). Twenty healthy (American Society of Anaesthesiologists, physical status Class 1), non-smoking, male volunteers with a BMI of 18-30 kg/m^2^ and aged between 18 – 45 took part in the study. All subjects gave informed consent, with experimental procedures approved by the UK National Research Ethics Service in South East Wales. Subjects were screened for personal history of neurological or psychiatric disease by the Mini International Neuropsychiatric Interview (Sheehan et al., 1998). Exclusion criteria further included contraindications to MEG or magnetic resonance imaging (MRI), and self-reported needle phobia.

### Visual paradigm

In order to induce gamma oscillations in the visual cortex (Swettenham et al., 2009), subjects were presented with an annular, stationary, square-wave visual grating of spatial frequency 3 cycles per degree on a mean luminance background. Gratings were displayed 150 times, with 75 at 100% contrast and 75 at 75% contrast (Muthukumaraswamy and Singh, 2013). Grating visual angle was 8º. Subjects were instructed to focus on a small, red, continually displayed dot at the centre of the grating. To maintain attention, participants were asked to press a button with their right index finger at stimulus offset. Gratings were displayed for 1.5 to 2 s (jittered offset), with a 3 s inter-stimulus interval. Gratings were displayed on a Sanyo XP41 LCD back-projection system displaying at 1024×768 at 60 Hz.

### MEG recordings, MRI and analyses

The 275-channel MEG suite at CUBRIC (CTF MEG by MEG International Services) was operated in the supine position for comfort and to minimise head movements during sedation. A further 29 channels were recorded for noise cancellation. Fiducial (reference) coils were attached 1 cm superior to the nasion and, bilaterally, 1 cm anterior to the tragus for assessment of head position within the MEG dewar. MEG data were sampled at 1200 Hz using axial gradiometers analysed as synthetic third-order gradiometers (Robinson and Vrba, 1999; Vrba, 2001).

All subjects underwent a structural T1-weighted scan, for co-registration to MEG, using a GE HDx 3T MR scanner with 8 channel head coil. A fast spoiled gradient echo sequence was obtained with 1 mm isotropic voxel resolution.

Pre-processing of MEG data included trial rejection by visual inspection for gross artefact using CTF DataEditor. Visual responses were localised using the synthetic aperture magnetometry (SAM) beamformer spatial filter with 4 mm isotropic resolution. A global covariance matrix (i.e. including all trials) was computed on data filtered to the visual gamma range, 30 – 80 Hz. Pseudo-*t* statistics were computed for baseline (−1.5 – 0 s) vs. stimulus (0 – 1.5 s). Virtual sensors were reconstructed in visual cortex at the location of peak *t*-statistic on an individual dataset basis. This grating paradigm induces peaks bilaterally in visual cortex. Spectra were computed using a fast Fourier transform of the virtual sensor.

### Ketamine and placebo infusion protocol

All subjects underwent both a ketamine and a placebo infusion, in counterbalanced, pseudo-random order, with at least 2 weeks between sessions. The ketamine infusion consisted of 0.5mg/kg (of body mass) racemic ketamine hydrochloride in 50 ml saline. For placebo, 50 ml saline only was used. Infusions were administered by intravenous cannula in the dorsum of the left hand, with an initial bolus (∼ 1 minute) of 0.25 mg/kg followed by maintenance infusion of .25 mg/kg delivered over 40 minutes. Maintenance infusion was controlled by an Asena-PK infusion pump (Alaris Medical, UK).

### Modelling analysis

In direct comparison with the default, canonical 4-population NMDA model (cmm_NMDA.m), we implemented a novel thalamo-cortical model based upon the same Morris-Lecar (1981) conductance equations, as per Moran and colleagues (Gilbert et al., 2016; Moran et al., 2011c; Muthukumaraswamy et al., 2015). The model consisted of 6 inter-connected cortical populations (pyramidal, interneuron and stellate) and 2 thalamic populations (reticular and relay), as depicted in figure 1. Populations were ‘connected’ in a hierarchical format (figure 1) and in accordance with prior descriptions (Moran et al., 2011c), however, the model does not explicitly encode the spatial distribution of populations or receptors.

**Figure 1.**
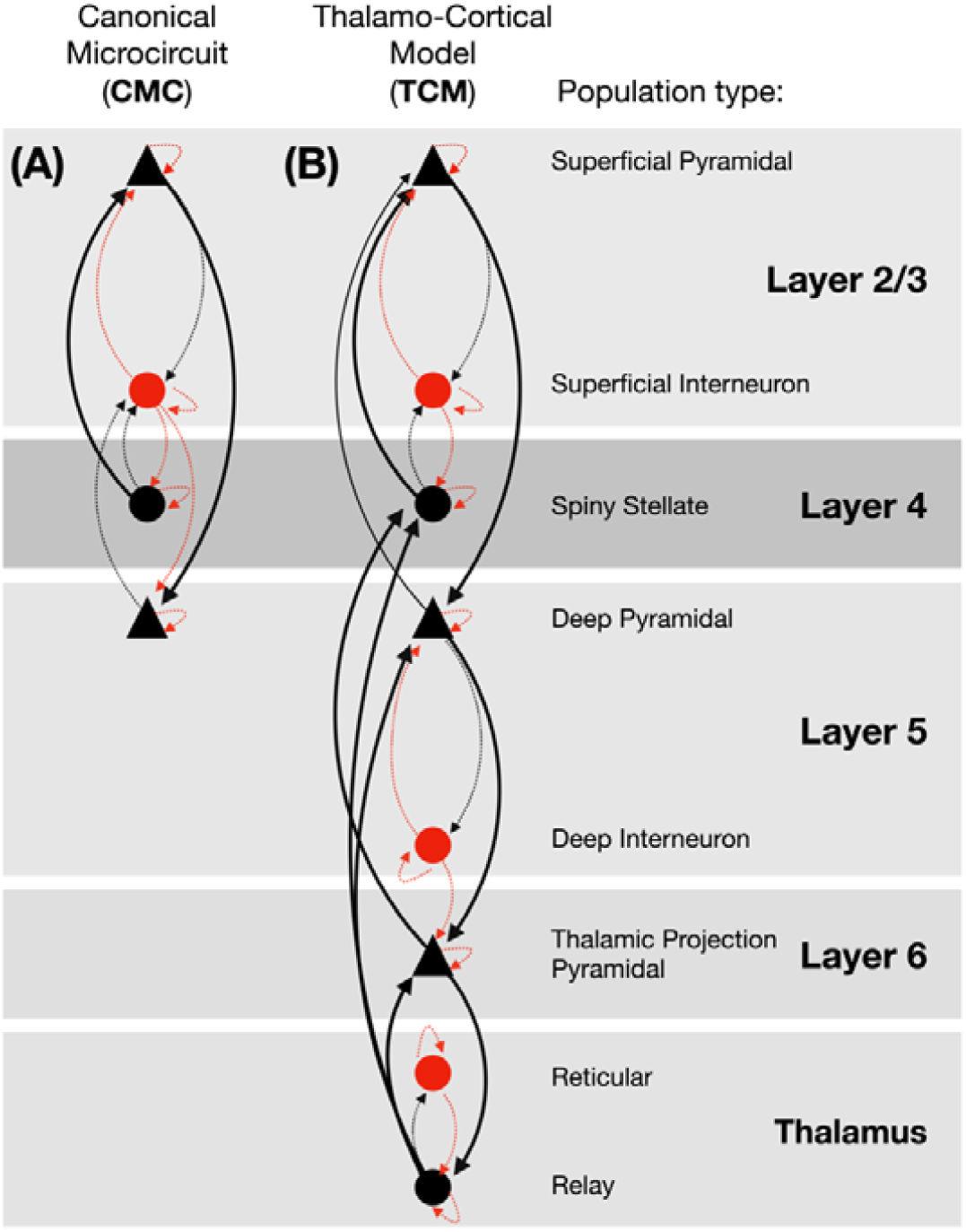
Schematic of the architecture and connectivity of the conductance based 4-population canonical microcircuit (CMC) and the novel 8-population thalamocortical model (TCM).

The model architecture comprised pyramidal and interneuron populations in layer 2/3 and layer 5, a stellate population in layer 4, and a thalamic-projection pyramidal population in layer 6. The time evolution of each population was governed by coupled differential equations describing the rate of change of membrane potential and rate of change of conductance supported by particular receptors;

Where V is the membrane potential, *g*_*n*_ is the rate of change of conductance conferred from receptor *n* and *g*_*n*_(*V* -*V*_*n*_) represents the contribution to the membrane potential of channel n, given its reversal potential (*V*_*n*_). C is the membrane capacitance and *u* is any exogenous or latent endogenous input current.

In the conductance rate equation, *k*_*n*_ represents the time constant, or decay rate, of currents mediated by channel *n*. Conductance, *ς*_*n*_, is calculated from the third term in equation 1, where the coupling parameter, *y*_*ij*_, coupling population *j* to *i*, is multiplied by the expected firing of source population *j*. The sigmoid function sigma represents the cumulative distribution function of the presynaptic depolarisation around a threshold potential of V_R_ = −40 mV, which determines the proportion of cells firing (Marreiros et al., 2009; Moran et al., 2011c). A full explanation of the parameterisation of these equations for model fitting is included in the supplementary materials.

Conductance terms are included for 6 channels: AMPA, NMDA, GABA_A_, GABA_B_, m-currents and h-currents, from voltage-gated and leak potassium channels respectively. Only layer 6 thalamic-projection pyramidal cells and thalamic relay populations possess m and h channels (see figure 1). This formulation is an extension of the NMDA model of Moran (Gilbert et al., 2016; Moran et al., 2011c) (CMM_NMDA in the SPM DCM toolbox) and includes the same voltage switch to represent the NMDA magnesium block;

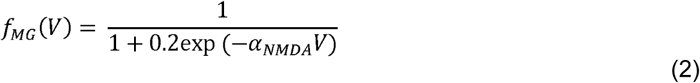

Reversal potentials and decay rates (full list in table S1) were obtained from existing models (Gilbert et al., 2016; Moran et al., 2011c) and from the literature (GABA_B_: Gerstner et al. (2014), M-channel: Bordas et al. (2015) and Brown and Adams (1980), H-channel: Gonzalez et al. (2015)).

M-channels are responsible for m-currents; non-inactivating voltage-gated potassium currents, which may be important to the intrinsic dynamics (oscillations) of the cell membrane (Bordas et al., 2015). They play a role in setting inhibition-excitation balance (Peters et al., 2005) and are unusual in that they are found open at rest (Nicholls et al., 2012). H-channels, which produce the cardiac funny current, produce hyperpolarisation activated currents, which not only have a role in the regulation of the resting membrane potential (Santoro and Baram, 2003) but may contribute to macroscopic network oscillations (Gonzalez et al., 2015). The G-protein coupled GABA_B_ receptor has also been linked to macroscale oscillations, including in the gamma range (Brown et al., 2007), possibly through its ability to re-balance the excitation-inhibition balance through interactions with NMDA receptor mediated currents (Gandal et al., 2012). As such, we parameterise all of these model elements in order to explore whether they may be altered under ketamine.

Both the CMC and TCM models were integrated numerically over a 2 s period (ensuring steady-state) at dt = 1/1200 s using an Euler method with delays, as per David et al (2006). This differs to standard DCM for spectral densities (DCM-CSD and DCM-SSR), since, by default, these methods do not use a numerical integration in the time domain. Instead, making some assumptions about steady-state, a transfer function computes a set of kernels using the system’s eigenspectrum, which shape parameterised output noise. In the present study, we opted instead to perform an explicit integration, using a scheme similar to that implemented in DCM for ERPs. This Euler method incorporates delays using the scheme:

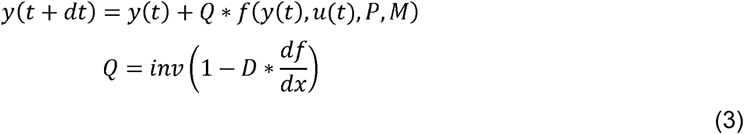

Here, y(t) is a vector of model states, u(t) is the model input (time invariant, D.C.), P and M are data structures containing the model parameters and function handles, respectively. The delay operator, Q, in the update scheme is computed from the number-of-states by number-of-states delay matrix and the system Jacobian, df/dx (as per DCM for ERPs). For the CMC model there are 20 states (4 populations each with 5 states [mV, gAMPA, gGABA-A, gNMDA and gGABA-B] – here we added gGABA-B). For the TCM model, there are 56 states (8 populations each with 7 states, [mV, gAMPA, gGABA-A, gNMDA, gGABA-B, gM and gH]). Note that while we added GABA-B channels to the default cortical model, we did not add M- or H-channels, since these were only present on L6 pyramidal cells and thalamic relay populations, which do not exist within the cortical model. Differentiation for df/dx was computed using the finite difference method implemented in spm_diff.m.

This method differed from the DCM for ERP approach, since we drive the model with a constant/ direct current into thalamic relay populations, as opposed to a Gaussian bump-function. This change was motived by the empirical demonstration that sustained, induced oscillations in visual cortex continue for as long as a static visual grating is displayed (Koelewijn et al., 2013).

Dynamic mode decomposition (DMD) (Schmid, 2010) was applied to the resulting population-by-time membrane potentials of contributing cells (SS, SP and DP in the CMC and SS, SP, DP & TP in the TCM), resulting in a set of frequency-specific temporal modes (Brunton et al., 2016; Schmid, 2010). A frequency-domain output was computed using a smoothed FFT (Matlab) of this mode series (see Supplementary Methods). The use of DMD was motivated by the observation that each population within the model could produce a complex oscillatory output (i.e. more than a single frequency). Because these frequencies overlap between populations, using a simple weighted sum of the population timeseries resulted in an additional level of non-linearity in the models during fitting (because the parameter derivative at a given frequency could be mediated by a number of underlying states/populations generating signal at that specific frequency). Thus, we employed DMD (which is a variant of principal component analysis specifically for dynamical systems (Schmid, 2010)) as a mechanism for identifying the principal oscillations present in the LFP.

In the TCM, the cortex-thalamus-cortex loop incorporated a delay of 11 ms, made up of 8 ms cortex to thalamus and 3 ms thalamus to cortex, in accordance with previous models (Lumer, 1997; Traub, 2004). However, since there is large variation in modelling estimates of these delays in the literature (Adams et al., 2020; Hashemi et al., 2017; Roberts and Robinson, 2008) we parameterised these delays, allowing them to vary during data fitting.

The models were fitted to the MEG virtual sensor frequency spectrum between 4 and 80 Hz using the DCM Gauss-Newton optimisation routine (a Variational Laplace / Bayesian inversion routine for nonlinear models, see spm_nlsi_GN.m), which minimises a free energy term (Friston et al., 2007, 2006; Penny, 2012) in order to maximise the model evidence and produce a-posteriori estimates for the model parameters, which included intrinsic connectivity parameters (*γ* in eq.3) and receptor time constants for AMPA, GABA_A_, NMDA, GABA_B_, m-channels and h-channels (*κ* in eq.1). The DCM optimisation routine rests on a gradient ascent optimisation, where the objective function (minimising free energy) is similar to a complexity-adjusted, precision-weighted sum of squared error (SSE) term.

Fitting was performed in two stages for each model (the 4-population model and TCM): first, a mean dataset was generated from the 32 datasets and inverted to find a set of priors from which to fit individual datasets. Second, all 32 datasets (16 subjects, 2 models each: 1x ketamine, 1x placebo) were fit with full models, starting from the same priors. We have previously demonstrated that this approach is sensitive to individual differences in model parameters induced by pharmacological manipulation (Shaw et al., 2017). Furthermore, it does not require an a-priori selection of which parameters may explain differences between the drug conditions, as would be the case using a multi-condition single DCM per subject.

### Study objectives

In this study, we had 3 key objectives. First, to compare the evidence for a novel 8-population thalamo-cortical model against an existing 4-population cortical model, in terms of their free energy and using Bayesian Model Selection.

Second, to assess how well the model recapitulated the broadband spectrum of the MEG virtual sensors, and whether the model fits were adequately accurate to recapitulate the drug effects on the spectrum.

Third, given adequately fitted models, we were specifically interested in examining the effects of ketamine, as compared to placebo, on 2 sets of model parameters. The parameter sets included (1) the time-constants of the modelled receptors (AMPA, NMDA, GABA_A_ and GABA_B_, m- and h-channels) and (2) the synaptic connectivity between populations, as reflected in figure 1 and detailed in supplementary figure S4. Between drug differences were determined by paired-*t* test with 5000 permutations and omnibus correction for multiple comparisons (Nichols and Holmes, 2001).

## Results

### Participants

Of 20 recruited subjects (mean age 25.7, SD 6.2), 1 subject withdrew prior to ketamine infusion, and an error in MEG acquisition was made for another. A further 2 datasets from the visual paradigm reported here, were rejected based on visual inspection of gross recording artifacts (reported previously (Shaw et al., 2015)), resulting in 16 usable, complete (i.e. including ketamine + placebo) datasets for analysis.

### Bayesian Model Selection

Bayesian Model Selection (BMS) using a fixed-effects (FFX) model showed greater evidence for the TCM (figure 3, A). While we assume the same underlying model for all subjects, it is possible that the effects of ketamine could have induced changes outside the scope of the model, therefore suggesting that some subjects had (due to drug) an altered underlying model. As such, we also computed BMS using random effects (RFX). RFX BMS also showed TCM as the winner (TCM: expectation of the posterior = 0.97, protected exceedance probability = 1, Bayesian Omnibus Risk (BOR) = 7.68×10^−9^ Figure 3 B). The mean model fits for each model are shown in figure 3 C and D. Individual dataset model fits are included in the supplementary materials figure 3.

**Figure 2.**
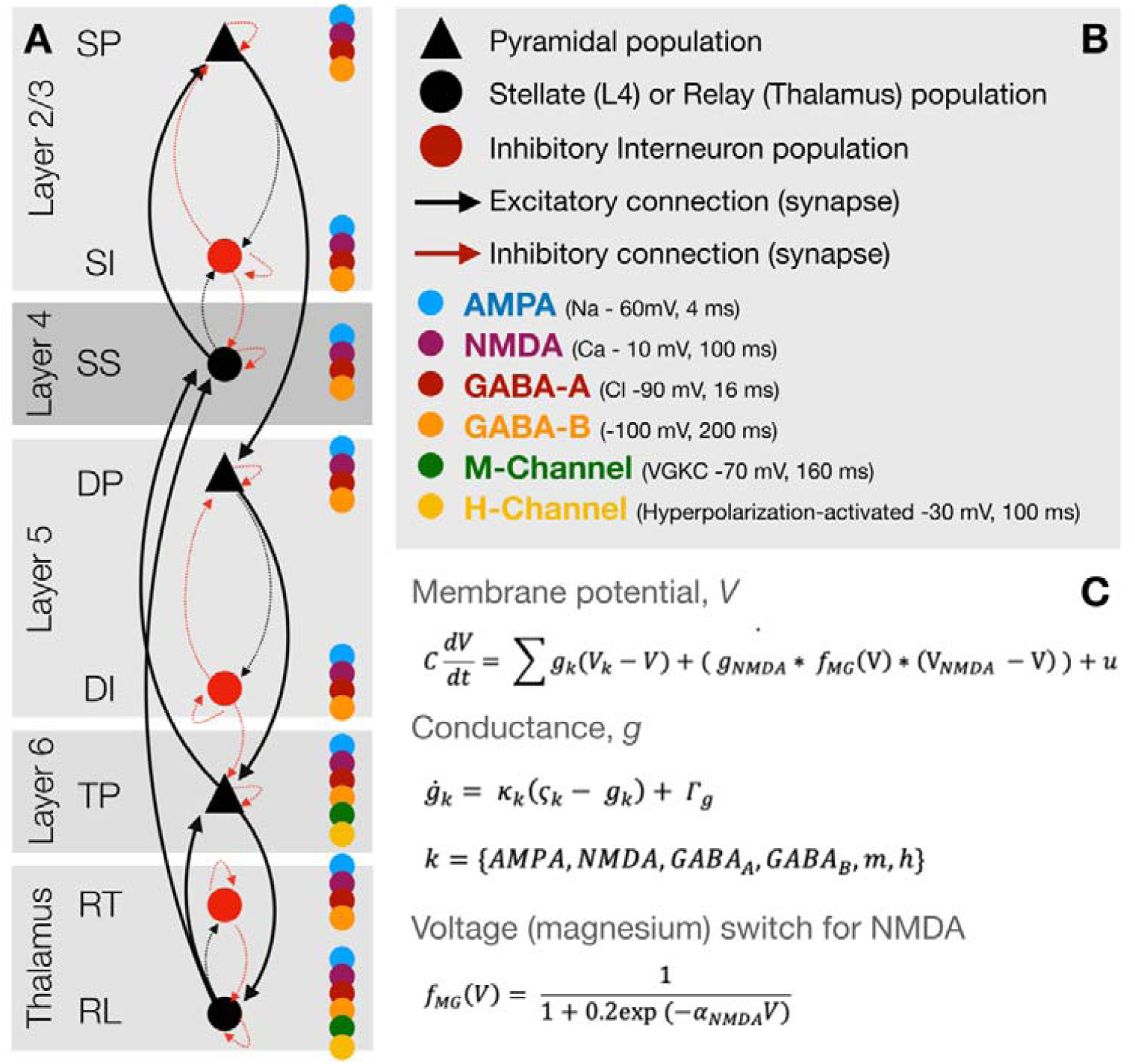
(A) Overview of the thalamo-cortical column architecture and connectivity. Coloured dots represent which channels are present on the population. (B) Key for the symbols in (A). (C) Summary of the conductance-based equations of motion for each population. SP = layer 2/3 (superficial) pyramidal, SI = superficial interneurons, SS = layer 4 spiny-stellates, DP = layer 5 (deep) pyramidal, DI = deep interneurons, TP = thalamic projection pyramidal (with M- and H-currents), RT = thalamic reticular and RL = thalamic relay.

**Figure 3.**
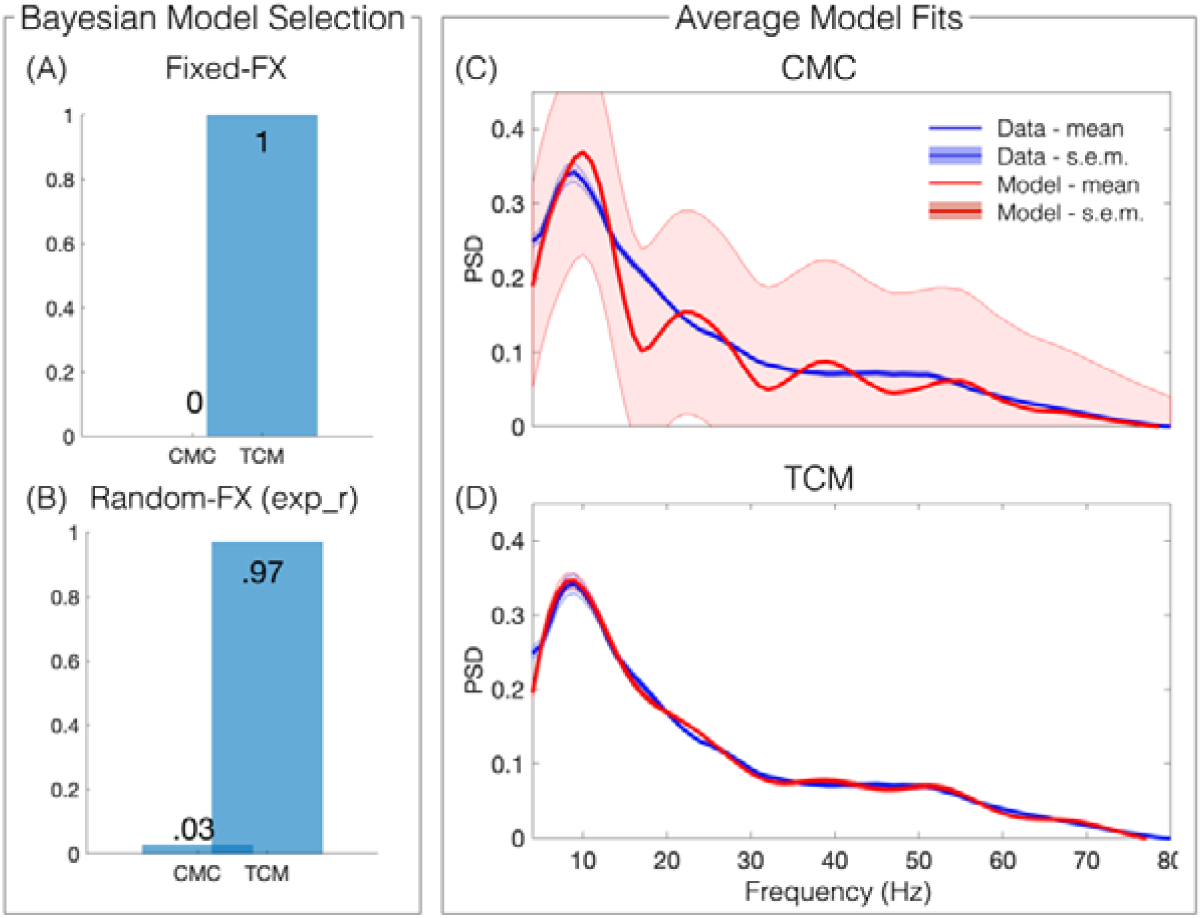
Bayesian Model Selection using Fixed- and Random-effects selected the TCM over the CMC as the winner. Averaged model fits (with standard error of the mean) show high accuracy of the spectral fits with the TCM model. Panel (A) demonstrates the posterior probabilities from the fixed-effects BMS, panel (B) shows the same for the random-effects BMS, which had an exceedance probability = 1 and Bayesian omnibus risk = 7.68×10^−9^. Panel (C) shows the mean and stand-error over datasets of the spectral density of the data (blue) and CMC-model fits (red). Panel (D) shows the same as (C) but for the extended, TCM-model.

### Model fits & recapitulating broadband spectral changes induced by ketamine

Although model fitting and subsequent analyses are carried out in the frequency domain, our numerical integration method allowed for visual confirmation of the presence of sustained membrane potential oscillations in all datasets. Assessment of the model fits (figure 4) demonstrated good correspondence between the MEG virtual sensor spectrum, and model predicted spectrum. Quantifying this using a correlation of the amplitude at each frequency point across subjects (figure 4, A) between the real and model spectrum revealed that, over the group, the model was able to recapitulate the whole spectrum, declining only at higher frequencies (> 70 Hz) and only non-significantly at >70 Hz. Furthermore, the free energy (FE) values for each dataset are depicted in figure 4 (C), clearly showing greater FE for the TCM in all datasets.

**Figure 4.**
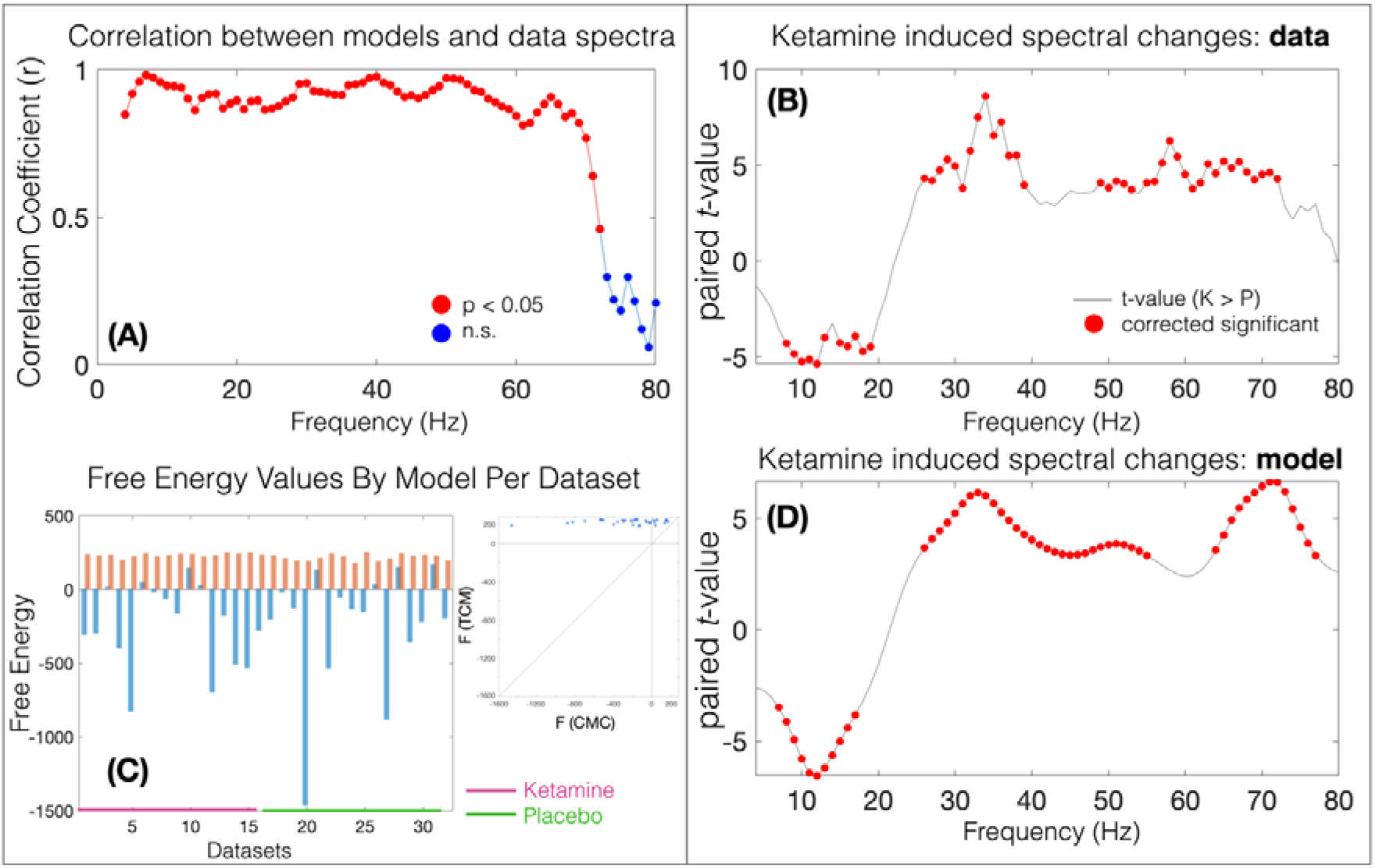
(A) The correlation of the amplitude at each frequency step between MEG virtual sensor (‘real data’) and model predictions. Red dots represent p<=0.05. (B) Red dots represent significant change in amplitude at this frequency point with ketamine in the MEG spectra; the amplitudes of low frequencies are attenuated by ketamine while amplitudes of high frequencies are increased by ketamine. (D) Showing the same drug-effects as (B) in the model predicted spectrum. (C) shows the free energy values for each dataset and for each model (red = TCM, blue = CMC).

We used a paired-t statistic with 5000 permutations and omnibus correction for multiple comparisons (Nichols and Holmes, 2001) to quantify ketamine induced changes in the spectra of both the MEG virtual sensor data, and the model predicted spectra (figure 4, B and D). In accordance with previous reports (Hong et al., 2010; Lazarewicz et al., 2010) and with previous analyses of this data (Shaw et al., 2015), ketamine reduced the amplitude of lower frequencies (theta, alpha) and increased the amplitude of higher, gamma frequencies (figure 2c). Applying the same procedure to the model output revealed the same statistical pattern, demonstrating that the model fits were sufficiently accurate enough to recapitulate the drug-induced spectral changes (figure 4 D).

### Model parameters analysis: receptor decay constants and intrinsic coupling

We employed randomisation based (5000 permutations) paired-t tests with omnibus correction to evaluate changes in the synaptic connectivity and receptor time constants. No significant changes were observed in receptor time constants (in equation 1) with drug (mean percent change [and standard deviation]: AMPA = −0.003 [.007], GABA_A_ = −0.001 [.003], NMDA = −0.49 [3.3], GABA_B_ = 0.001 [0.002], M = −12 [17.3], H = −8 [24]).

Applying the same randomisation-based statistical procedure to the 41 synaptic connectivity estimates (see supplementary figure 4 for connectivity schematics) revealed 4 highly significant parameter changes with ketamine compared to placebo. These consisted of 2 NMDA-mediated connections: SP←SI (t=30755, p<.001, mean change with ketamine +470%, SD = 0.56), DP←DP (t=8.5^+15^, p<.001, mean change with ketamine +933%, SD = 1.2). They further included the AMPA mediated SP←SI (t=7055, p<.001, mean change with ketamine +470%, SD = 0.56) and GABA_A_ mediated DP←DP (t=9.6^+15^, p<.001, mean change with ketamine +1000%, SD = 1.16). Figure 5 shows the individual dataset parameter changes for each of the 4 parameters.

**Figure 5.**
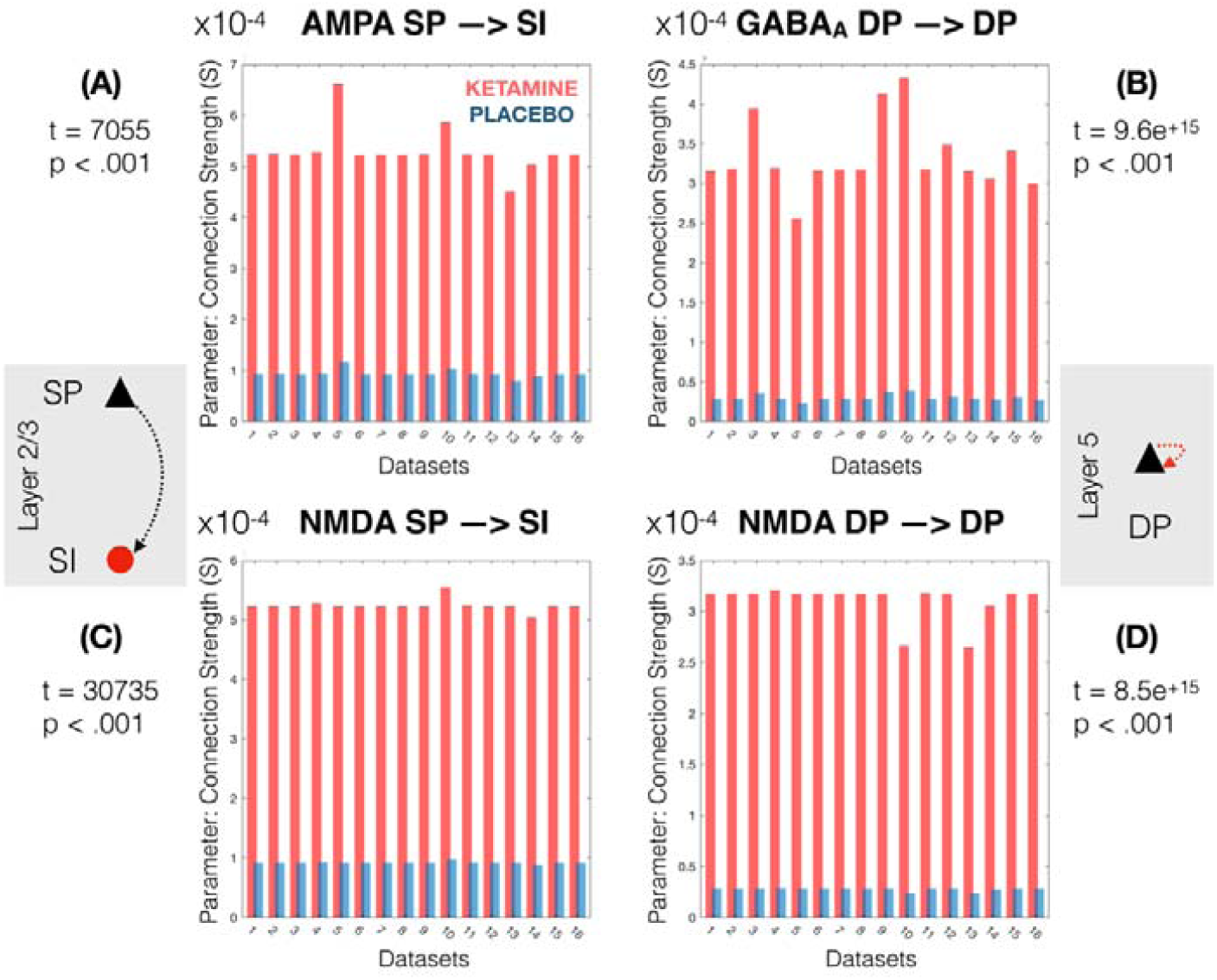
Showing individual dataset values for the 4 parameters significantly altered by ketamine. These include (A) increased AMPA mediated SP-→SI, increased NMDA mediated SP→SI, increased GABA_A_ mediated DP→DP and increased NMDA mediated DP→DP.

## Discussion

The current study demonstrated that a simplified thalamo-cortical model, implemented with DCM, is able to recapitulate the spectral changes induced by subanaesthetic ketamine administration during a visual grating task, showing good correspondence with MEG recorded changes. Furthermore, direct comparison with the existing NMDA model (cmm_NMDA.m) provided strong evidence in favour of the TCM. Parameter level effects revealed that compared to saline placebo, ketamine enhanced NMDA and AMPA mediated superficial pyramidal to superficial interneurons (SP→SI) and both NMDA and GABAA mediated self-gain of deep pyramidal populations (DP→DP).

### Comparing the TCM to the NMDA model: Bayesian Model Selection

Bayesian model selection comparing a 4-population cortex-only model with the novel thalamo-cortical model (TCM) suggested that the broadband, 4 – 80 Hz spectra obtained from a visual grating paradigm in MEG was better explained by the TCM. BMS with random effects (RFX) indicated very strong evidence in favour of the TCM (BOR=7.68×10^−9^). This is consistent with theories which propose a role for thalamo-cortical communication in the generation and pace-making of lower frequency oscillations (Halgren et al., 2017; Lumer, 1997). However, one interpretation of the BMS result could be that the additional dimensionality of the TCM model, which has 8 rather than 4 populations, simply permits a richer repertoire of responses which enable it to better explain individual or drug differences in the data spectrum. However, the free energy term used to compare the models accounts for this additional complexity (Friston and Penny, 2011; Penny, 2012; Stephan et al., 2009), suggesting that the TCM is better at explaining the data even when compensating for the additional complexity.

An alternative approach to explaining the alpha and gamma changes observed with ketamine could have been to run models separately to explain just the band limited changes. We chose not to adopt this approach, since the complexity and dimensionality of the TCM are designed to fit broadband spectra. As such, fitting only a limited frequency band would result in a redundancy within the model space (and parameters), leading to a lack of specificity in the model components.

Future studies fitting broadband spectral responses may wish to consider this model and integration scheme, particularly if their data features either violate assumptions about stationarity required for DCM-SSR/CSD, or if the research question requires access to the continuous time output of the system as well as frequency domain.

### Model sensitivity: spectral and parameter changes

Before making any between-drug parameter inference, it was important to assess whether the model had summarized the data well enough to be sensitive to the drug induced changes in the spectrum. As shown in figure 4 (sub-plots B/D), the spectral difference in amplitude observed in the empirical MEG data (4B), which included significant reductions in alpha amplitude and increases in gamma amplitude, were also evident at the same frequencies in the model output (4D). Without this assurance that the model had accurately recapitulated the data features, any parameter inference would be arbitrary.

Comparing the models fitted to ketamine and placebo, we identified 4 synaptic parameters that were significantly enhanced by ketamine. These connections included both NMDA and AMPA mediated superficial pyramidal to superficial interneurons (SP→SI) and both NMDA and GABA_A_ mediated self-gain of deep pyramidal populations (DP→DP). Our (model dependent, *in silico*) observation that AMPA and NMDA mediated SP→SI and NMDA mediated DP→DP are all increased by NMDA receptor (NMDAR) antagonism initially appears to be at odds with the idea of antagonising excitatory receptors. However, the paradox of NMDAR antagonism producing cortical excitation is established (Homayoun and Moghaddam, 2007; Jackson et al., 2004). One explanation for this is that NMDAR hypofunction reduces excitation of GABAergic interneurons, leading to disinhibition of pyramidal cells (Homayoun and Moghaddam, 2007). Our parameter increases in superficial layer SP→SI (AMPA and NMDA mediated) may therefore reflect this increased state of excitability – linking increased pyramidal population NMDA and AMPA excitability with the downstream effects of increased gamma amplitude and decreased alpha amplitude.

A previous study which used a cortical DCM model to explore ketamine induced changes in network (rather than local) connectivity also reported increased excitatory connectivity under ketamine (Gilbert et al., 2018). In this study, the NMDA mediated forward connection from somatosensory (S1) to right frontal cortex was enhanced by ketamine in healthy subjects. Taken together, our findings suggest that ketamine may increase both forward extrinsic and intrinsic NMDA mediated connectivity.

However, the increase in both GABA_A_ (inhibitory) and NMDA (excitatory) within-population connectivity in layer 5 deep pyramidal cells is less interpretable by the disinhibition explanation. Although these parameters both reflect local (within population) mechanisms on the same population, the GABA_A_ parameter mediates local inhibition, while the NMDA parameter mediates local excitation. Increases in both of these parameters concomitantly with ketamine suggests that the local dynamics of layer 5 pyramidal cells are significantly, and perhaps non-linearly, altered by NMDAR antagonism.

We suggested that ketamine’s ability to attenuate lower frequency oscillations might reflect alterations in thalamo-cortical connectivity, since alpha oscillations in visual cortex are coupled to the firing of visual lateral geniculate nuclei (Lorincz et al., 2009). Moreover, other anaesthetic agents, such as propofol, alter cortico-thalamic and thalamo-cortical dynamics (Hashemi et al., 2017). However, our results did not reveal any ketamine-induced changes either in the thalamo-cortical or cortico-thalamic delays, nor connectivity measures, even though BMS clearly showed that the TCM was a better model of the data than the CMC. Our results therefore suggest that, while broadband spectra from visual cortex are better explained by a thalamo-cortical model, the spectral changes induced by ketamine may mostly reflect ketamine’s effects in cortex.

### Study limitations and future work

Given the sensitivity of model parameters to manipulation by ketamine, it would be interesting to know whether any of the observed effect are dose-dependent, particularly since ketamine has seen a resurgence in popularity due to its antidepressant effects. In the present study, we used a fixed 0.5mg/kg dose, however there have been no human neuroimaging-based studies of ketamine at varying levels, which would permit discovery of dose-dependent effects, such as has been done in rodent using DCM (Moran et al., 2014).

This study explored the effects of ketamine in healthy, young participants. However, the same modelling framework could be applied to clinical trial data, where ketamine is currently under investigation as a treatment option for a variety of psychiatric (mood) disorders. In this context, the sensitivity of model parameters could serve as sensitive and specific biomarkers – both of illness and of treatment response.

Our sample was limited to young (mean age = 26, sd = 6) males. Such a homogenous sample may not be representative of a more heterogenous population. Moreover, we had only 16 complete (i.e. ketamine and placebo) and useable datasets. In a previous analysis of the change in gamma with ketamine, we *post-hoc* estimated a large effect size of 1.78 (Cohen’s D). However, this estimate was drawn from the same data as the present analysis, not from an independent sample.

In this study, we have demonstrated that a thalamo-cortical circuit, fitted to empirical pharmaco-MEG data with DCM better explains broadband spectra from visual cortex than a reduced 4-population cortex-only model. We further show that this TCM can accurately and significantly recapitulate spectral changes induced by ketamine and is sensitive to synaptic connectivity dynamics. We have shown that enhanced cortical connectivity mediated by NMDA, AMPA and GABA_A_ plays a role in shaping the oscillatory responses of ketamine, not just NMDA. Finally, our results suggest that the oscillatory changes observed with ketamine are mediated by changes in cortex rather than in thalamo-cortical or cortico-thalamic connectivity. Our results should be considered by subsequent imaging-based investigations of ketamine, and extended to studies of patient groups undergoing ketamine therapy, where elucidating the mechanism of action of ketamine is of crucial importance.

## Model population abbreviations

SP: Superficial layer pyramidal (L2/3)
SI: Superficial layer interneuron (L2/3)
SS: Spiny Stellate (Layer 4)
DP: Deep layer pyramidal (Layer 5)
DI: Deep layer interneuron (Layer 5)
TP: Thalamic projection pyramidal (Layer 6)
RT: Thalamic Reticular
RL: Thalamic Relay

## Acknowledgements

Designed research: ADS, SDM, KDS

Performed research: ADS, SDM, NS, KDS

Development of model: ADS, NA, RLS, RJM, KDS

Analysed data: ADS

Wrote the paper: ADS, SDM, NS, NA, RLS, RJM, KDS

## Data & Code Availability

The matlab code for the thalamo-cortical model and integration functions, in a DCM-compatible format, will be made available via GitHub (http://github.com/alexandershaw4/) upon publication of this manuscript. The modelling data that support the findings of this study are available from the corresponding author, upon reasonable request.

