## supplementary materials for "Generative modelling of the thalamo-cortical circuit mechanisms underlying the neurophysiological effects of ketamine"

**Supplementary Materials: Thalamo-cortical oscillatory model for ketamine – Shaw et al 2019**

Parameterisation of the neural and observation models.

Here we aim to set out how the model equations were parameterised in order to invert the model to the empirical ERPs. Elements of the equations which are parameterised are highlighted in bold. Starting with the conductance equation:

$$\varsigma_{n}=\gamma_{i,j}\sigma(\mu_{v}^{j}-V_{R}, \sum j)$$

Eq a.

Gamma is a (parameterised) sparse matrix of connectivity parameters between subpopulations, such that $\gamma_{i,j}$, describes the synaptic strength of the connection from population j to population i. This matrix (which, if full, would be 8x8) is the parameter of primary interest as the estimated parameters correspond to the effective (synaptic) connectivity between subpopulations.

We include one such matrix for AMPA and GABA-A/GABA-B mediated connectivity (where excitatory elements are AMPA and inhibitory, including diagonals, are GABA-A/B), and a separate matrix for NMDA. This extension of the cmm_NMDA code allows us to separately model AMPA and NMDA excitatory local connectivity with minimal change to the equations.

Lower case sigma, $\sigma$, represents the expected (average) firing rate of the source population j (sigma is a vector of 8 parameters, each corresponding to the firing rate of one of the populations). V_R_ is a fixed threshold of -40 mV for firing (not a parameter).

The second component of the conductance equation:

$$\dot{g}_{n}= \kappa_{n}\left( \varsigma_{n}- g_{n} \right)$$

Eq b.

This update scheme describes the evolution of the conductance, determined by the rate constant of channel n ($\kappa_{n})$ multiplied by the change in conductance (the output of equation a) minus the same quantity from the last time step. The rate constant (K) is a parameterised vector of length 6 (1 value each for AMPA, NMDA, GABA-A, GABA-B, M- and H- channels). These receptor ‘rates’ are common across all subpopulations.

The computed conductances for each channel (g_n_) permit calculation of the voltage equation (again, an update-scheme):

$$\frac{\mathrm{dV}}{\mathrm{dt}}=(\sum g_{n}\left( V-V_{n} \right)+u)/C$$

Eq c.

Here, V is the membrane voltage of the population. V_n_ is the reversal potential of channel n (fixed, not a parameter) and u is any endogenous or exogenous input current (a single parameter, in this experiment a D.C / invariant in time, which enters only thalamic relay cells). C is the membrane capacitance (parameter vector of length 8; one capacitance for each subpopulation).

Numerical integration of equation c leads to a membrane-potential (voltage) time series for each population. In DCM, this time-series is referred to as a hidden state, since it is not directly observed in the empirical data, rather an observation (aka forward) model is applied to this timeseries in order to compare it to the empirical MEG data.

Observation model.

The purpose of the observation model is to transform the integrated channel-by-time voltage series into a local field potential (LFP)-like spectrum, for comparison to the MEG virtual sensor spectrum. In this study we assume that the MEG virtual sensor data is a weighted sum of the voltage series of the principal (pyramidal) cells. Due to the complex oscillatory nature of the membrane series, where the timeseries of each population demonstrates oscillations of multiple frequencies, we employed Dynamic Mode Decomposition (DMD) to separate the contributing population timeseries into a set of frequency-specific modes (DMD is introduced in (Schmid, 2010) and has been used for neural data previously, for example in (Brunton et al., 2016)). The single-channel output spectrum is computed by parameterised (n=4) weighted sum of the smoothed Fourier transformed mode data. Finally, an electrode gain parameter, L, was applied. Supplementary figure 1 summarises the integration and observation steps and supplementary figure 2 illustrates the DMD routine.

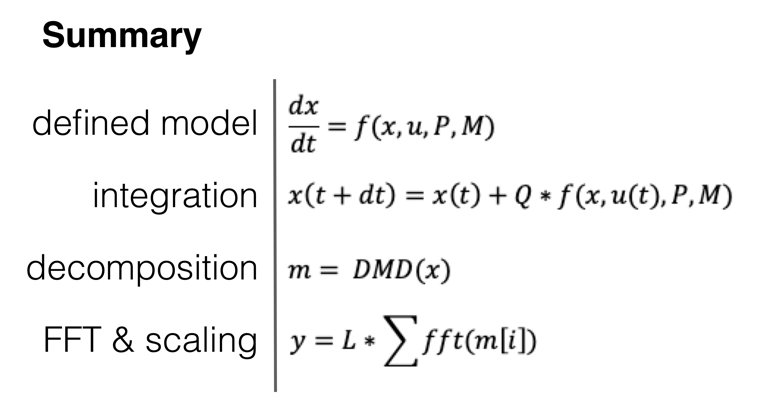

Supplementary Figure S1. Summary of the generative model specification, integration, decomposition and spectral output steps.

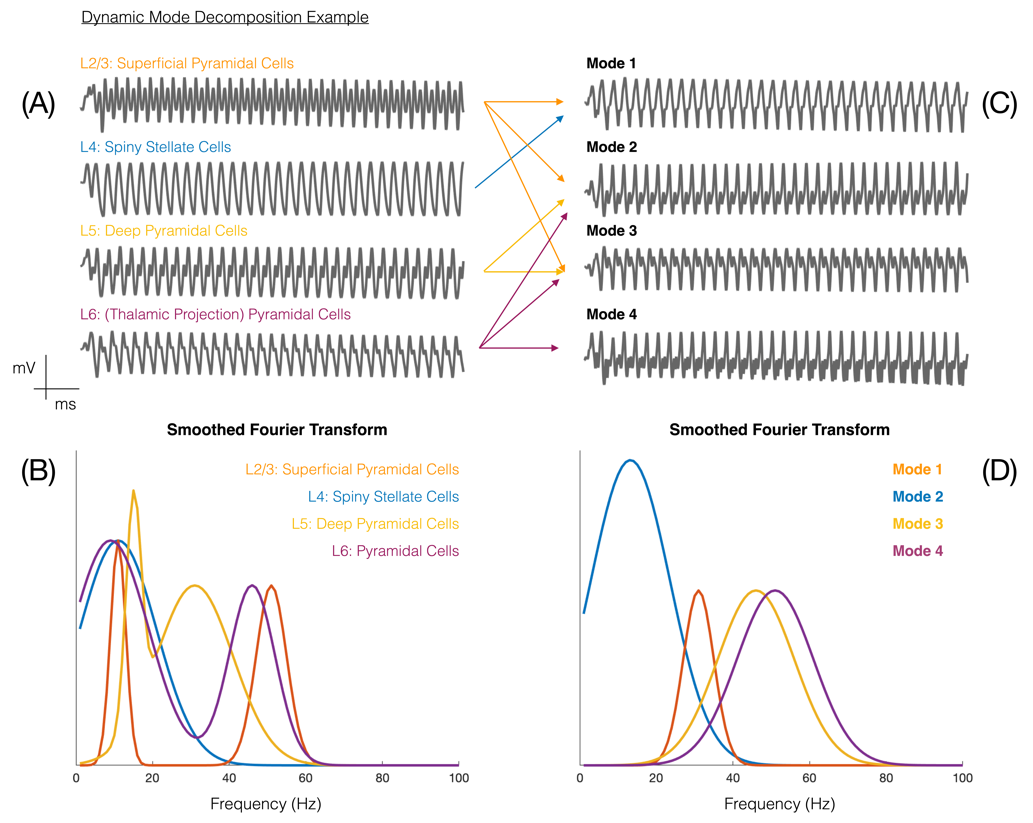

Supplementary Figure 2. Depicts the use of dynamic model decomposition (DMD) to compute a set of frequency modes from the complex oscillatory signals of the contributing populations membrane potentials. (B) shows a smoothed Fourier transform of each of the 4 contributing cell populations (A), where each population displays multiple oscillatory frequencies. We employed DMD to decompose these complex oscillations in the time domain into 4 discrete frequency modes (C), each of which has a weight parameter in the frequency domain (D).

| Channel | ion | Reversal (mV) | Decay rate (ms) |
| --- | --- | --- | --- |
| Potassium Leak | K | -70 | ~ |
| AMPA | Na | 60 | 4 |
| NMDA | Ca | 10 | 100 |
| GABAA | Cl | -90 | 16 |
| GABAB | ~ | -100 | 200 |
| M | K | -70 | 160 |
| H | Non  selective | -30 | 100 |

Supplementary Table S1. Reversal potentials and receptor decay constants for each channel in the model.

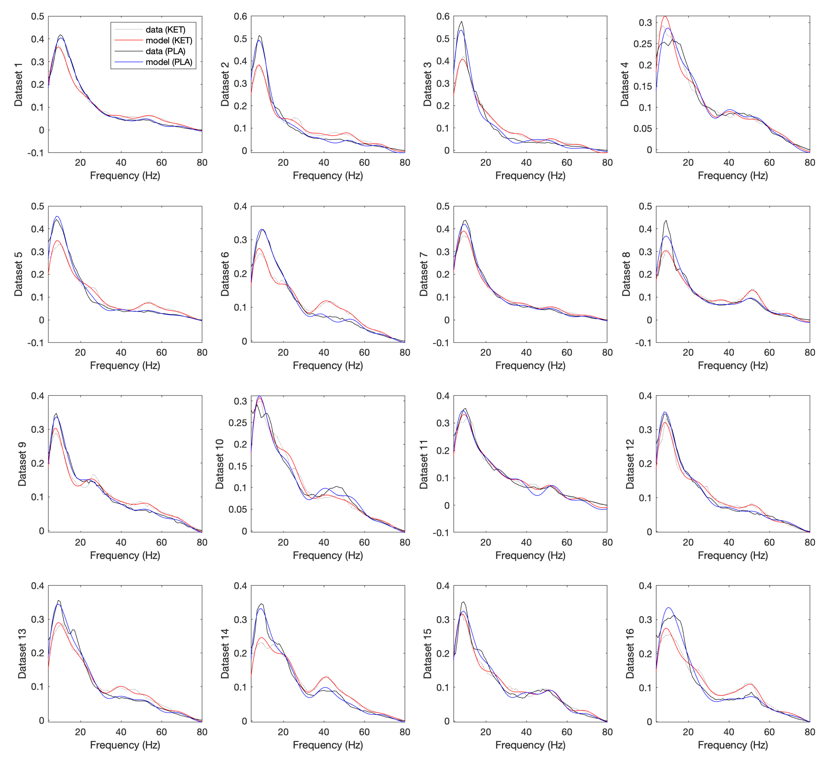

Supplementary Figure S3. Individual dataset model fits.

| **Parameter** | **Description** | **Prior Mean** | **Variance** |
| --- | --- | --- | --- |
| H / Hn (41 parameters) (see Sup. Fig. 4) | Intrinsic connection strengths | Excitatory = 4  Inhibitory = 8 | 1/8 |
| TC | Log scale parameter on Receptor Rates in table 1 | 0 | 1/8 |
| Input (u) | D.C to Relay population | 2 | 1/8 |
| Delays (D) | Thal🡪Cort Cort🡪Thal | 3 ms [0] 8ms [0] | 1/8 |
| Weights [J] | Weightings on frequency-modes (x4) | 1 | 1/8 |
| Electrode Gain [L] | Scales output spectrum | 1 | 1/8 |

Supplementary Table S2. Parameters with non-zero variances – i.e. which varied during model fitting.

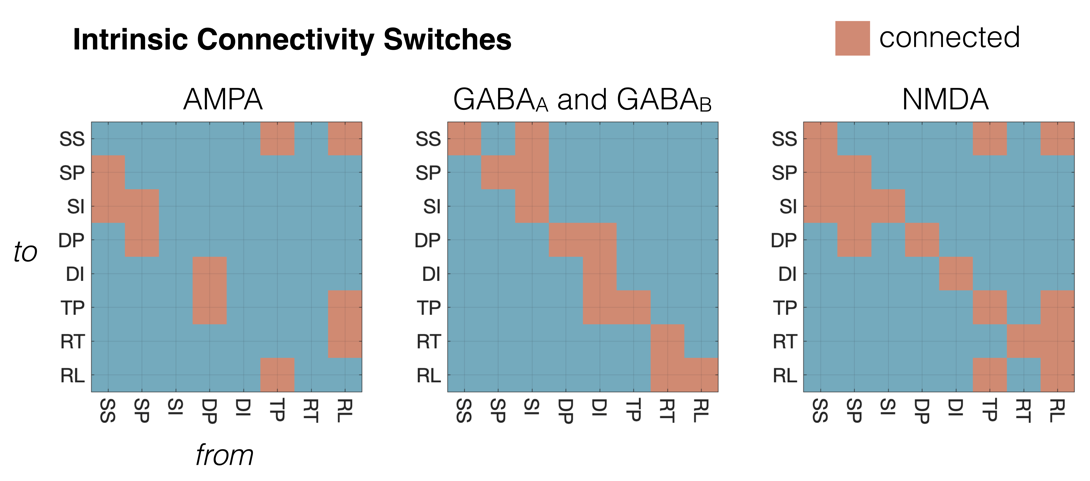

Supplementary Figure S4. Intrinsic Connectivity Switches for AMPA, NMDA and GABA-A/B. Population abbreviations: **SS** = L4 Spiny Stellates, **SP** = L2/3 (Superficial Layer) Pyramidal Cells, **SI** = (Superficial Layer) Inhibitory Interneurons, **DP** = L5 (Deep Layer) Pyramidal Cells, **DI** = L5 (Deep Layer) Inhibitory Interneurons, **TP** = L6 (Thalamic Projection) Pyramidal Cells, **RT** = Thalamic Reticular Cells, **RL** = Thalamic Relay Cells.
